# Genomic foundation model embeddings encode higher-order viral genome architecture beyond sequence composition: a benchmark of Evo 2

**DOI:** 10.64898/2026.07.14.738542

**Authors:** Deyvid Amgarten, Anderson Schinaid, Fernanda de Mello Malta, Alexandre R. Marra, João Renato Rebello Pinho

## Abstract

Genomic foundation models such as Evo 2 are increasingly applied to microbial genomics, yet how well their representations capture viral genome organisation, and how reliably they generate viral sequence, remain poorly characterised. We present a reproducible benchmark of Evo 2 on viral genomes. Using a pre-registered RefSeq viral corpus (19,429 genomes, organised by Baltimore class and host domain), we evaluated three axes: linear probes decoding Baltimore class, host domain and viral family from mean-pooled embeddings; ridge-regression probes recovering genomic features, including higher-order architectural properties such as gene density, coding fraction and gene overlap; and generative completion of fragmented genomes, scored on a leakage-safe set of eukaryote-infecting viruses (excluded from Evo 2’s training corpus by design) against a bacteriophage comparator. All probes used cross-validation with sequence-identity–clustered folds, benchmarked against both a GC-and-length control and a 6-mer composition representation. From its optimal intermediate layer, the 20B embedding classified Baltimore class at 0.96 accuracy and host domain at 0.99, exceeding both baselines; for viral family, however, 6-mer composition (0.89) matched the embedding (0.91. Most informatively, the embedding decoded coding fraction, gene density and gene overlap (R² = 0.61, 0.77 and 0.64) far beyond 6-mer composition (0.10, 0.38 and 0.27), evidencing genuine encoding of genome architecture rather than nucleotide composition (p < 0.001). Performance scaled with model size. In generation, perplexity was lower for bacteriophages (1.18 bits/nt) than for held-out eukaryotic viruses (1.80). Evo 2 encodes functional viral genome architecture beyond composition, while taxonomic and generative behaviour partly reflect composition and training exposure.

## 1. Introduction

Foundation models based on *Large Language Model* (LLM) architectures have transformed computational biology by extending self-supervised learning to DNA sequences at genome scale. Training on large genomic datasets containing trillions of nucleotides has enabled these models to learn transferable representations for diverse downstream tasks, including variant effect prediction, functional annotation, regulatory element identification, and biological sequence design (Nguyen et al., 2024; Brixi et al., 2025). Among these models, Evo 2 represents a major advance by combining large-scale autoregressive modeling with long-context processing, enabling the capture of biological dependencies spanning from the molecular level to complete genomes (Brixi et al., 2025).

The rapid evolution of genomic language models has been accompanied by substantial changes in sequence representation strategies. DNABERT-2 replaced fixed-length *k*-mer tokenization with *Byte Pair Encoding* (BPE), reducing computational redundancy while improving learning efficiency across multiple species (Zhou et al., 2023). More recently, architectures such as Caduceus have incorporated biological priors directly into the model by enforcing reverse-complement invariance, allowing biologically equivalent DNA strands to share consistent representations (Schiff et al., 2024). In parallel, an increasing number of genomic foundation models have adopted auxiliary tokens and conditioning mechanisms, including *conditioning tokens*, *species tokens*, and *strand tokens,* to provide explicit biological context during training or inference. These mechanisms enable models to incorporate external biological information without modifying the nucleotide sequence itself, improving contextual awareness across heterogeneous genomic domains (Nguyen et al., 2024; Sanabria et al., 2024; Veiner & Supek, 2026).

Despite these architectural advances, the biological information encoded within latent representations remains only partially understood. Most previous studies have focused on developing improved architectures, tokenization strategies, or conditioning mechanisms, whereas comparatively few have systematically investigated which biological properties are intrinsically recoverable from model embeddings (Alain and Bengio, 2016). This question is particularly relevant for viral genomes, whose compact organization, high evolutionary diversity, and extensive structural heterogeneity provide a challenging benchmark for evaluating representation quality.

Viral genomes are an unusually stringent test bed for such representations, because their architecture varies widely and largely independently of nucleotide composition (Koonin et al., 2020). The virosphere is polyphyletic, with replication modules and structural genes repeatedly recombined over evolutionary time, and the seven Baltimore classes that formalise this diversity are properties of the replication cycle rather than of the deposited sequence: a foundation model reads every class as the same four-letter DNA alphabet. Physical organisation is equally extreme. In the corpus assembled here, genome or segment length spans four orders of magnitude — from a few hundred base pairs to 2.5 Mb — GC content ranges from 18% to 79%, and coding density covers its entire attainable range, from small viruses that compress their genomes through overlapping reading frames to large DNA viruses rich in non-coding content. Because these architectural properties vary largely orthogonally to composition, viral genomes allow a probe to separate learned genome organisation from mere nucleotide statistics.

In this work, we systematically evaluate the latent representations produced by Evo 2 (20B Base) using Ridge regression-based linear probing, without modifying the underlying architecture or introducing additional conditioning mechanisms. We assess the extent to which the learned embeddings encode biologically relevant properties of viral genomes, including Wright’s Effective Number of Codons ENC (Wright, 1990), GC content, CpG dinucleotide frequency, gene density, and coding fraction, all computed from coding sequences (CDSs) extracted from official GenBank annotations. In addition, we benchmark both the representational and generative capabilities of the model using a controlled evaluation framework that distinguishes information genuinely captured by the embeddings from signals attributable to simple sequence composition, providing a reproducible protocol for assessing genomic foundation models in viral genomics.

## 2. Materials and Methods

### 2.1 Viral genome corpus

We assembled a viral genome corpus from the NCBI RefSeq viral release (O’Leary et al., 2016, release of May 2026; 19,429 genome records, including individual segments of segmented viruses). Each record was mapped to a Baltimore replication class and a host domain (eukaryote, bacterium, archaeon) by joining family- and genus-level taxonomy to the International Committee on Taxonomy of Viruses (ICTV) Virus Metadata Resource (VMR, MSL41) (Lefkowitz et al., 2018). Per-genome genomic features (length, GC content, coding fraction, gene density, gene and CDS counts, non-coding and intergenic statistics) were computed directly from the GenBank flat-file annotation. Quota groups (Baltimore class × host) and quality cut-offs were fixed in a versioned pre-registration file prior to analysis to avoid post-hoc selection (Nosek et al., 2018) (Supplementary Table S4). For each probe we drew balanced subsets from this corpus (details below). No deduplication was applied at corpus-assembly stage: RefSeq viral is curated at approximately one reference genome per species, and redundancy among the genomes actually drawn into each probe was instead assessed and controlled directly via sequence-identity clustering at the cross-validation stage.

### 2.2 Genomic feature definitions

Architectural features were computed per record directly from the RefSeq GenBank flat files (.gbff) using Biopython. Coding base pairs were taken as the length of the union of all CDS exon intervals (join-aware, so that alternatively spliced or multi-exon CDSs are counted once), and coding fraction as coding base pairs divided by genome length (bounded in [0, 1]); non-coding content as genome length minus coding base pairs. Gene density was defined as the number of annotated gene features per kilobase of genome (falling back to the CDS count when a record lacks explicit gene features). Mean intergenic length was the average gap between consecutive, non-overlapping CDS loci. Gene overlap (overlap_bp) was quantified at the locus level as the summed length of all CDS loci minus the length of their union — that is, the total base pairs covered by more than one CDS — a hallmark of genome compression in small viruses, and was log-transformed for probing. GC content was computed over the full genome. As deliberate low-order composition controls, CpG and UpA dinucleotide depletion were computed as observed-to-expected ratios from each full genome sequence, O/E_XY = f(XY) / [f(X)·f(Y)], with f(XY) the observed relative dinucleotide frequency and f(X), f(Y) the mononucleotide frequencies (UpA computed as TpA on the DNA strand; genomes with fewer than 100 resolved ACGT bases were excluded from these statistics). Records lacking an inline sequence (CON-type entries) were skipped at extraction.

### 2.3 Model and embedding extraction

We benchmarked the Evo 2 20B base model (configuration evo2_20b, 1M-token context) as the primary model (Brixi et al., 2025), with the 7B base model (evo2_7b) as a smaller-scale comparator; neither was fine-tuned. The 20B model was run on an NVIDIA H100 80GB GPU (Hopper architecture), whose native support for FP8 input projections matches the precision for which the checkpoint is calibrated; the 7B model was run on an NVIDIA L40S (48 GB), a non-Hopper GPU on which FP8 input projections were disabled so that all layers ran in bfloat16 (∼13 GB resident). Per-genome embeddings were mean-pooled over tokens; for the 20B model we report the intermediate layer blocks.18.mlp.l3, selected by the layer-sensitivity analysis of Section 2.5, and for the 7B model the layer blocks.28.mlp.l3. Genomes up to the context window were embedded in a single forward pass; longer genomes were processed in 32,768-bp windows with a 16,384-bp stride (capped per genome), and the resulting window embeddings were averaged, yielding one embedding vector per genome (8,192 dimensions for the 20B model, 4,096 for the 7B).

### 2.4 Representation probes

Representations were standardized and used as input to linear probes under repeated stratified 5-fold cross-validation (three repeats; 15 estimates per metric), reporting mean ± standard deviation. Classification of Baltimore class, host domain and viral family used multinomial logistic regression (balanced class weights, accuracy); decoding of continuous features used ridge regression with internal alpha selection (R²). Heavy-tailed features were log-transformed. Each probe was compared against two baselines computed on the identical genomes and folds: a trivial GC-and-length control (log-length and GC content) and a 6-mer composition representation (4,096 relative hexamer frequencies, matching the embedding dimensionality), which is a strong composition-only model. We tested whether the embedding outperformed each baseline with paired t-tests across folds. The classification subset for Baltimore comprised 150 genomes per class (n = 981 across seven classes); host and family probes used the feature subset (120 genomes per quota group; n = 1,200, all host domains; family restricted to families with at least 25 members, n = 349). The feature-regression subset was the same 1,200-genome set. To verify that scores were not inflated by near-identical genomes shared across cross-validation folds, we clustered every probed genome by sequence identity (MMseqs2 linclust (Steinegger and Söding, 2018); 95% identity, 85% coverage, greedy-set-cover mode — the same parameters pre-registered for corpus redundancy control) and repeated each probe under group-aware cross-validation (StratifiedGroupKFold for classification, a grouped analogue of k-fold for regression; three repeats, clusters as groups), so that no cluster could appear in both the training and held-out folds of a split.

### 2.5 Model scale and layer-sensitivity analysis

To test whether representation quality scales with model size, and to identify which embedding layer best captures viral genome structure, we embedded the same 1,912-genome probe subset with Evo 2 20B across five candidate layers (base, no fine-tuning; 24 hyena/attention blocks (Poli et al., 2023) and an 8,192-dimensional hidden state, versus 32 blocks and 4,096 dimensions for the 7B model). Embeddings were extracted on a single NVIDIA H100 80GB GPU (Hopper architecture), which, unlike the L40S used for the 7B model, natively supports FP8 input projections. Because the optimal embedding layer is not known a priori for an untested model, we extracted five candidate layers in a single forward pass per genome window — blocks.{11,15,18,21,23}.mlp.l3, spanning roughly 46–96% of network depth — using forward hooks that mean-pooled each layer’s output over tokens and transferred it to host memory immediately; this keeps the memory cost of extracting five layers comparable to that of extracting one. Windowing, striding and pooling otherwise followed the protocol described in Section 2.3. To assess whether the FP8/bfloat16 precision difference between the two GPUs confounds this comparison, a subset of layers was additionally extracted with FP8 disabled on the H100 (forcing bfloat16). This precision-matched extraction performed markedly worse than the native FP8 extraction across every target — most severely on fine-grained functional features (Supplementary Table S1) — indicating that the 20B checkpoint is calibrated for FP8 inference rather than precision-agnostic; a strict precision-controlled comparison was therefore not meaningful, and each model is instead reported in its native operating configuration (FP8 for the 20B on Hopper, bfloat16 for the 7B on a non-Hopper GPU). All five 20B layers were evaluated with the same probe battery, baselines, and cluster-aware cross-validation as the 7B model (Section 2.4; probed genomes and clusters are identical across models), and we report the full depth curve for every target — rather than only the best-performing layer — to avoid selection bias in the scale comparison. Because the 20B embedding is higher-dimensional than the 7B (8,192 vs. 4,096 components), which could by itself inflate linear-probe performance independently of representation quality, every probe was additionally re-run with both embeddings projected to a common, conservatively low dimensionality (150 components — set below the smallest per-fold training-set size across all targets) via PCA fit inside each cross-validation fold to prevent leakage.

### 2.6 Generative evaluation

Generative capability was assessed on a leakage-safe held-out set of eukaryote-infecting viruses (n = 100; excluded from OpenGenome2 by design, Brixi et al., 2025) and a comparator set of bacteriophages (n = 100; included in training), each requiring at least 8,192 bp. We computed two quantities. Perplexity was measured as teacher-forced bits per nucleotide over the first 8,192 bp of each genome (mean of the negative base-2 log-probability of the next nucleotide). Genome completion simulated a recovered contig: the first 4,096 bp served as a prompt, from which the model generated the following 1,024 bp (temperature 0.7, top-k 4) (Fan et al., 2018); the generated segment was compared with the true downstream sequence by 4-mer spectrum cosine similarity, and the model’s teacher-forced bits per nucleotide on the true gap quantified predictive fit. A 4th-order Markov model fitted on the prompts provided a composition baseline for both metrics. Because Evo 2 is autoregressive, completion uses only upstream context, matching the realistic task of extending a contig 5′→3′.

### 2.7 Reproducibility

The pre-registered corpus design, the feature-extraction and composition code, and the analysis notebooks (with embedded outputs) are released in the project repository; figures were produced by python deterministic scripts from the cached embeddings (available at the project repository). The corpus derives from public NCBI RefSeq and the ICTV VMR. Analyses used scikit-learn (Pedregosa et al., 2011). The layer-sensitivity sweep, scale comparison and dimensionality-matched PCA control (sweep_layers_20b.py, scale_analysis.py, pca_control.py) and the sequence-identity cluster map (cl95_cluster.tsv) used for group-aware cross-validation are released alongside the probe and figure-generation scripts.

## 3. Results

### 3.1 Genome type and taxonomy are strongly encoded

Linear probes recovered viral genome type and taxonomy from the Evo 2 20B embeddings (Figures 1A and 1B, Table 1). Baltimore class was classified at 0.961 ± 0.011 accuracy across seven classes, significantly above both the 6-mer composition baseline (0.808 ± 0.023) and the GC-and-length control (0.468 ± 0.026; both p < 0.001), and above the 7B model (0.883 ± 0.025; Section 3.4). Reverse-transcribing (VI, VII) and dsDNA classes were classified most accurately, whereas the single-stranded RNA classes were hardest, with most confusion occurring among them (Figure 1A). Host domain was classified at 0.995 ± 0.005, again above 6-mer (0.934 ± 0.018) and GC-and-length (0.816 ± 0.019). For viral family, however, the 6-mer composition baseline (0.891 ± 0.025) matched the embedding (0.907 ± 0.032; difference not significant), even though both far surpassed GC-and-length (0.354 ± 0.049). Thus genome type and host are encoded by the embedding beyond what composition alone provides, whereas family-level taxonomy is essentially a compositional signature that a simple k-mer model captures equally well. Sequence-identity clustering showed the probed genomes to be almost entirely non-redundant (only 16 of 1,912 merged at 95% identity, reflecting RefSeq’s one-reference-per-species design), and group-aware cross-validation changed every metric by at most 0.012—within the fold-to-fold standard deviation—confirming that these scores are not inflated by near-duplicate leakage (Supplementary Table S2).

**Figure 1.**
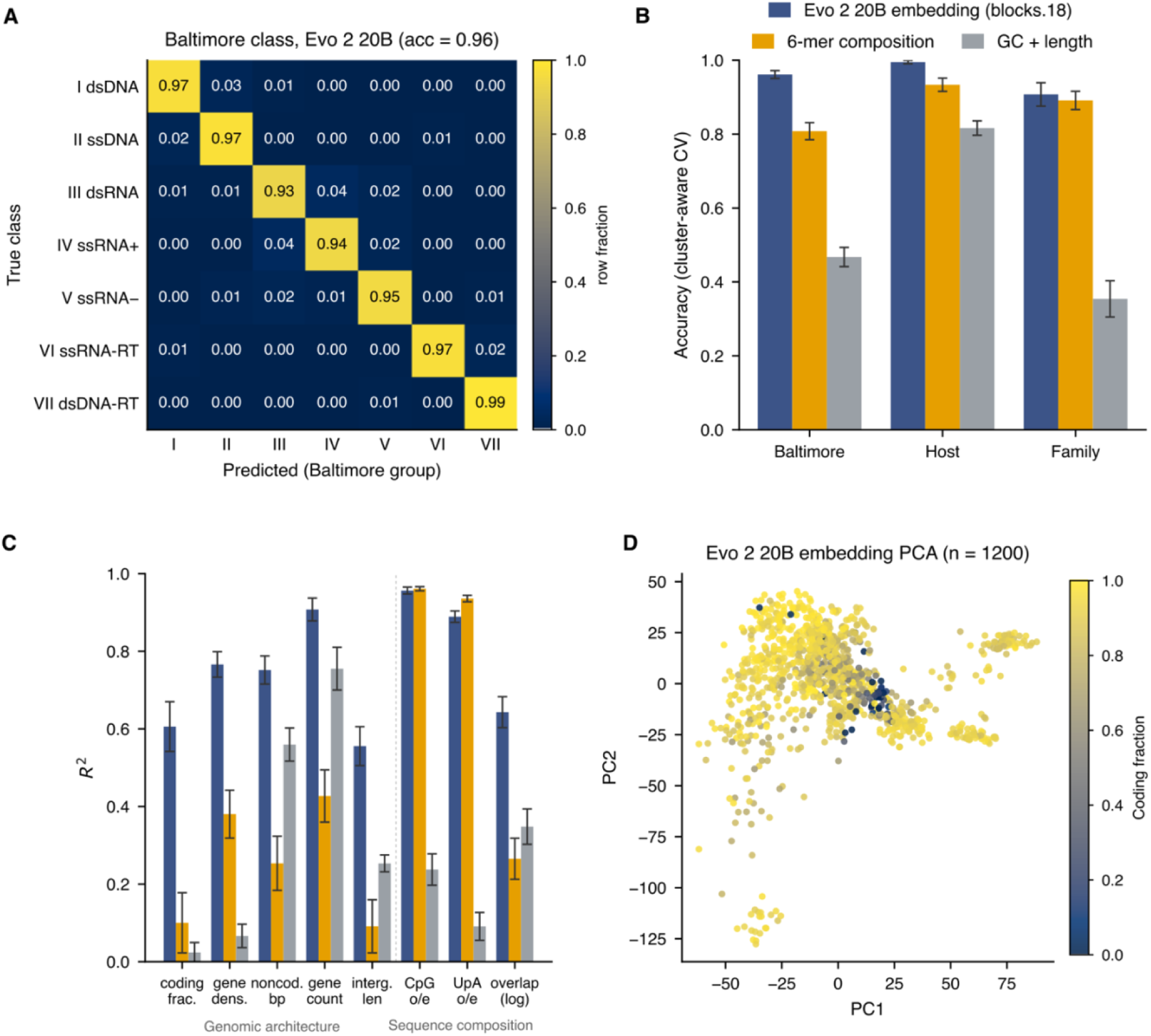
Representation probes for viral genome type, taxonomy and functional architecture (Evo 2 20B, layer blocks.18). (A) Row-normalized confusion matrix for Baltimore-class classification from the 20B embeddings (five-fold cross-validation; overall accuracy 0.96). (B) Classification accuracy (repeated cluster-aware cross-validation; mean ± SD) for Baltimore class, host domain and family, comparing the Evo 2 20B embedding with a 6-mer composition representation and a GC-and-length control. (C) Cross-validated R² per genomic feature (repeated cluster-aware CV; mean ± SD) for the same three representations. Features are grouped into higher-order genome architecture (coding fraction, gene density, non-coding content, gene count, intergenic length, gene overlap), where the embedding dominates, and low-order sequence composition (CpG and UpA dinucleotide depletion), where the 6-mer baseline is competitive by construction. (D) Two-dimensional principal-component projection of the feature-subset embeddings (n = 1,200), coloured by coding fraction.

**Table 1.** Probe performance for the Evo 2 20B embedding (layer blocks.18), repeated cluster-aware cross-validation, mean ± SD. Classification reports accuracy; regression reports R². The embedding is compared with a 6-mer composition representation (4,096 features) and a GC-and-length control; the last two columns give the paired-t significance of the embedding versus each baseline (*** p < 0.001; ns, not significant). The CpG, UpA and gene-overlap regressions used random cross-validation, and per-fold significance was not computed for them (see text).

| Probe | Target | n | Evo 2 20B embedding | 6-mer composition | GC + length | vs k-mer | vs GC+len |
| --- | --- | --- | --- | --- | --- | --- | --- |
| Class. (acc) | Baltimore | 981 | $0.961 \pm 0.011$ | $0.808 \pm 0.023$ | $0.468 \pm 0.026$ | *** | *** |
| Class. (acc) | Host domain | 1080 | $0.995 \pm 0.005$ | $0.934 \pm 0.018$ | $0.816 \pm 0.019$ | *** | *** |
| Class. (acc) | Family | 349 | $0.907 \pm 0.032$ | $0.891 \pm 0.025$ | $0.354 \pm 0.049$ | ns | *** |
| Regr. ($R^2$) | coding_fraction | 1200 | $0.606 \pm 0.064$ | $0.100 \pm 0.078$ | $0.024 \pm 0.026$ | *** | *** |
| Regr. (R<sup>2</sup>) | gene_density | 1200 | 0.766 ± 0.033 | 0.380 ± 0.062 | 0.067 ± 0.030 | *** | *** |
| Regr. (R<sup>2</sup>) | noncoding_bp (log) | 1200 | 0.752 ± 0.036 | 0.254 ± 0.070 | 0.560 ± 0.043 | *** | *** |
| Regr. (R<sup>2</sup>) | n_genes (log) | 1200 | 0.908 ± 0.029 | 0.427 ± 0.067 | 0.755 ± 0.055 | *** | *** |
| Regr. (R<sup>2</sup>) | mean_intergenic_len (log) | 1200 | 0.556 ± 0.050 | 0.091 ± 0.069 | 0.253 ± 0.022 | *** | *** |
| Regr. (R<sup>2</sup>) | CpG o/e | 1200 | 0.957 ± 0.009 | 0.961 ± 0.005 | 0.238 ± 0.040 | — | — |
| Regr. (R<sup>2</sup>) | UpA o/e | 1200 | 0.889 ± 0.015 | 0.936 ± 0.008 | 0.091 ± 0.036 | — | — |
| Regr. (R<sup>2</sup>) | gene overlap (log) | 1200 | 0.643 ± 0.040 | 0.265 ± 0.053 | 0.348 ± 0.046 | — | — |

Note: for viral family the 6-mer baseline matches the 20B embedding (difference not significant).
Among the added features, only gene overlap favours the embedding; the CpG and UpA dinucleotide controls are, as expected, captured at least as well by the 6-mer baseline (see text).

### 3.2 Functional gene structure is encoded beyond composition

The strongest evidence that the embedding captures genome organization, rather than composition, came from feature regression (Figure 1C, Table 1). Coding fraction and gene density were almost entirely missed by both baselines—6-mer composition reached R² of only 0.100 ± 0.078 and 0.380 ± 0.062, and GC-and-length only 0.024 and 0.067—yet the 20B embedding decoded them at 0.606 ± 0.064 and 0.766 ± 0.033 (all p < 0.001). Because a 6-mer model encodes local composition (including codon-level signal) and still fails here, the embedding must encode higher-order gene organization. The same pattern held for gene overlap—a hallmark of genome compression in small viruses—which the embedding recovered (R² = 0.643 ± 0.040) far better than 6-mer composition (0.265 ± 0.053) or GC-and-length (0.348 ± 0.046). As a deliberate negative control, two low-order composition statistics—CpG and UpA dinucleotide depletion—were, as expected, captured at least as well by the 6-mer baseline (CpG 0.961 vs 0.957; UpA 0.936 vs 0.889), confirming that the embedding’s advantage is specific to higher-order architecture rather than to composition in general. For features partly determined by genome size (gene count, non-coding base pairs), GC-and-length already explained much of the variance, but the embedding remained the best predictor (n_genes R² = 0.908 vs 0.755 and 0.427; non-coding bp 0.752 vs 0.560 and 0.254). A principal-component projection of the embeddings shows organization consistent with coding density (Figure 1D).

### 3.3 Generative capability reflects training exposure

Generative metrics distinguished sequence the model had seen from sequence it had not (Figure 2). Genome perplexity was lower for bacteriophages, which were included in training (1.18 bits/nt), than for the held-out eukaryotic viruses (1.80 bits/nt), consistent with their exclusion from OpenGenome2. In the completion task, Evo 2 predicted the true downstream sequence with lower bits per nucleotide than the Markov baseline for both groups (eukaryotic viruses, 1.78 vs 1.97; bacteriophages, 1.15 vs 1.98), the margin being far larger for the seen bacteriophages. For free generation, the composition of the generated segment (4-mer cosine similarity to the true sequence) exceeded the Markov baseline for bacteriophages (0.79 vs 0.68) but not for the held-out eukaryotic viruses (0.71 vs 0.74). Thus, while Evo 2 assigns the true sequence higher likelihood than a composition baseline even out of distribution, the compositional fidelity of its sampled output on unseen eukaryotic viruses did not surpass that baseline.

**Figure 2.**
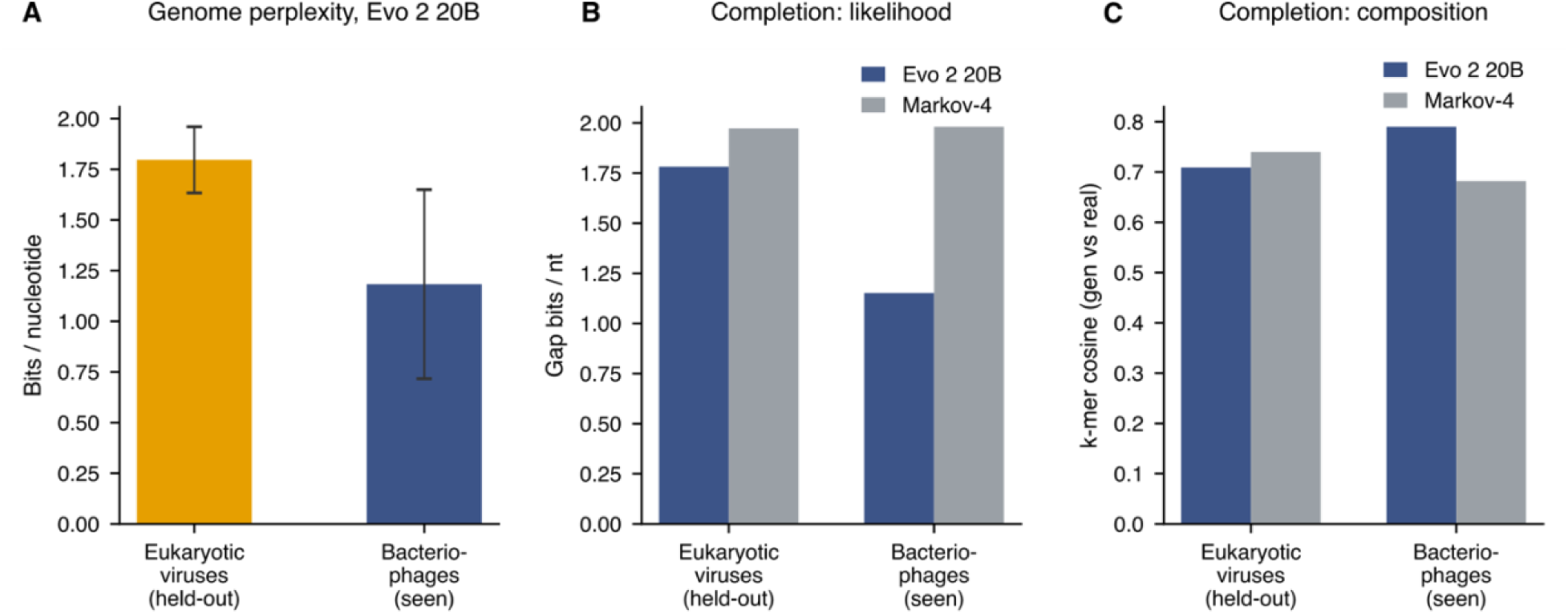
Generative evaluation for Evo 2 20B on held-out eukaryotic viruses versus seen bacteriophages. (A) Teacher-forced genome perplexity (bits per nucleotide; lower is better; bars show mean ± s.d.). (B) Completion likelihood on the held-out 1,024-bp gap for Evo 2 versus a 4th-order Markov baseline. (C) Compositional similarity (4-mer cosine) between the generated and true gap for Evo 2 versus the Markov baseline.

### 3.4 Representation quality scales with model size and depends on layer

The choice of layer was itself consequential: intermediate layers (blocks.11–18) carried the most linearly decodable structure, a mid-network layer (blocks.18) was the best single choice across targets, and the deepest layer tested (blocks.23) collapsed on every target — consistent with late layers specializing toward next-token prediction rather than a general-purpose genome representation (Figure 3A). We therefore report blocks.18 throughout, and provide the full depth curve for every target to make the layer dependence explicit rather than a hidden hyperparameter. The 20B model’s advantage over the 7B was systematic rather than incidental. Across the eight probe targets, the 20B embedding equalled or exceeded the 7B on the great majority — markedly so for Baltimore class (0.961 vs 0.883), gene density (0.766 vs 0.691), gene count (0.908 vs 0.825) and mean intergenic length (0.556 vs 0.367) — while remaining comparable on coding fraction and family (Figure 3B). Because the 20B embedding is higher-dimensional than the 7B (8,192 vs 4,096 components), which could by itself inflate linear-probe scores, we re-ran every probe with both embeddings projected to a common, conservatively low dimensionality (150 PCA components fit within each cross-validation fold); the 20B advantage persisted, indicating a genuine effect of model scale rather than of embedding width (Figure 3B, Supplementary Table S3).

**Figure 3.**
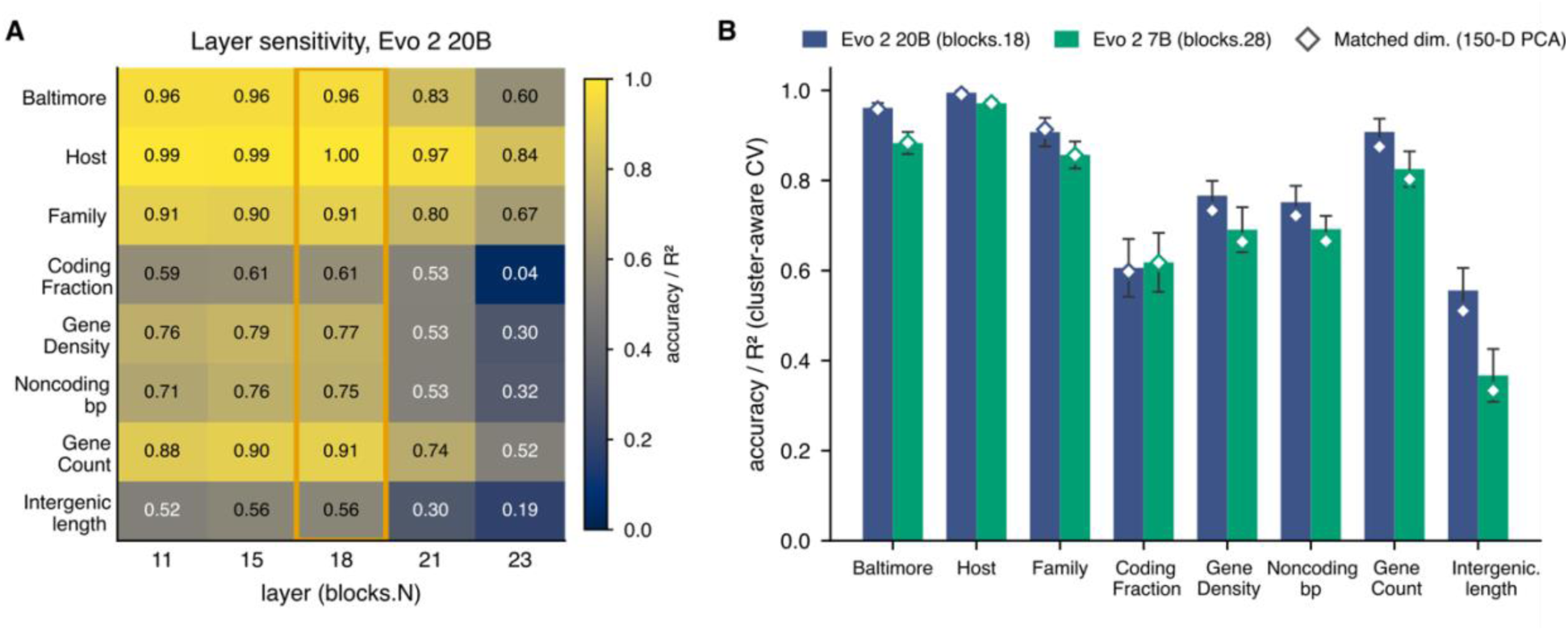
Representation quality by model scale and network depth (Evo 2, cluster-aware cross-validation). (A) Layer sensitivity of the Evo 2 20B embedding: cross-validated probe performance (accuracy for classification, R² for regression) for each target across the five extracted layers (blocks.11, 15, 18, 21, 23), spanning roughly 46–96% of network depth. Intermediate layers carry the most decodable structure; blocks.18 (highlighted) is the best single layer across targets, whereas the deepest layer (blocks.23) collapses. (B) Scale comparison of the Evo 2 20B (blocks.18) and 7B (blocks.28) embeddings per target (bars, mean ± SD); open diamonds show the same comparison after projecting both embeddings to a common 150-dimensional PCA subspace fit within each fold, confirming that the 20B advantage persists at matched dimensionality rather than reflecting embedding width. The same probe battery, baselines and cross-validation as in Table 1 were used.

## 4. Discussion

This systematic evaluation of Evo 2 (20B base) reveals that the model captures substantial biological structure in viral genomes beyond what simple sequence composition can explain. Across the three evaluation axes, Evo 2 demonstrates varying degrees of biological encoding, while comparison against a strong compositional baseline clarifies which information is genuinely contributed by the model beyond local sequence statistics. Benchmarking against a 6-mer representation, rather than relying solely on GC content and sequence length, demonstrates that the primary advantage of the learned embeddings lies in recovering higher-order aspects of genome organization. Coding fraction and gene density are accurately predicted from the embeddings (R² ≈ 0.61–0.77) but are only weakly explained by 6-mer composition (≤ 0.38), indicating that these properties cannot be reduced to local oligonucleotide frequencies. In contrast, viral family classification is explained at least as well by 6-mer composition, suggesting that family-level taxonomy largely reflects compositional signatures. Similarly, host domain remains partially predictable from sequence composition alone. The larger 20B model decoded these architectural features consistently better than the 7B, and the advantage survived matching both embeddings to a common dimensionality, indicating that representation quality genuinely improves with scale rather than merely with embedding width. This finding has practical implications for choosing model scale: the representational gains from larger models appear genuine rather than artifactual. The residual confusion among single-stranded RNA viruses highlights the current limitations of the learned representations and suggests that further gains may require domain-specific adaptation. From a representational perspective, these findings indicate that higher-order functional properties emerge naturally within the latent embedding space, contrasting with approaches that rely on auxiliary tokens or explicit conditioning mechanisms, where part of the biological context is provided directly as model input rather than inferred from sequence alone (Nguyen et al., 2024; Veiner & Supek, 2026).

The generative evaluation reveals a clear signature of training exposure. Both perplexity and sequence completion quality are consistently higher for bacteriophages, which were included in the OpenGenome2 training dataset, than for eukaryotic viruses, which were deliberately excluded from pre-training. Notably, even on these unseen viral genomes, Evo 2 assigns higher likelihood to the true sequence continuation than a Markov baseline, indicating that the model retains a degree of out-of-distribution generalization. However, freely generated sequences do not surpass the baseline in compositional fidelity, revealing a dissociation between probabilistic modeling and generation quality under distribution shift. This distinction is particularly relevant for applications such as metagenomic viral contig extension (Camargo et al., 2023), where producing biologically plausible sequence is more important than accurately ranking candidate continuations. In contrast to models that employ auxiliary conditioning or control tokens, Evo 2 relies primarily on information encoded during large-scale pre-training, making its generative performance more dependent on the quality of its internal representations than on externally supplied contextual information (Veiner & Supek, 2026).

Several limitations should be considered when interpreting these findings. Linear probes measure the decodability of information contained in the embeddings rather than its causal use during inference. In addition, the viral family classifier was restricted to families with sufficient representation, and although sequence identity-aware cross-validation produced essentially unchanged results, this likely reflects the relatively low redundancy of RefSeq viral genomes. More redundant datasets would require stricter controls to avoid overly optimistic performance estimates. The generative evaluation was also limited to a single prompt length, a fixed gap length, and moderate sample sizes, while autoregressive generation is known to deteriorate over longer sequence horizons. Furthermore, the leakage-free evaluation relies on the documented exclusion of eukaryotic viruses from the OpenGenome2 training dataset; direct verification against the complete training manifest would further strengthen this assumption. Finally, only the base Evo 2 model was evaluated. Future studies should investigate whether fine-tuning or explicit conditioning mechanisms improve robustness under distribution shift.

In conclusion, this work demonstrates that genomic foundation models can capture higher-order viral genome architecture beyond sequence composition, with representation quality scaling with model size and varying systematically across network depth. Overall, this work provides a reproducible benchmark for evaluating both the representational and generative capabilities of genomic foundation models on viral genomes and establishes a controlled evaluation framework that can support future comparisons across architectures, tokenization strategies, and conditioning paradigms.

## Conflict of Interest

The authors declare no other competing interests.

## Author Contributions

D.A. conceived and designed the study, implemented the corpus, probes and analyses, and wrote the manuscript. A.S. assisted with the generative-evaluation runs and contributed to results interpretation. F.M.M., A.R.M. and J.R.R.P. critically reviewed the study design and manuscript and contributed to interpretation of the results. All authors approved the final manuscript.

## Funding

This work was supported by the Genesis Genomics, which provided institutional research infrastructure. No specific grant was received for this project.

## Acknowledgments

The authors thank Genesis Genomics’ Information Technology and R&D teams and acknowledge cloud computing resources used for model inference.

## Data Availability Statement

The viral genome corpus derives from the public NCBI RefSeq viral release and the ICTV Virus Metadata Resource (MSL41). The pre-registered corpus design, analysis code and notebooks (including the figure-generation script) are available in the project repository: https://github.com/omicsintellab/benchmark-evo2-viral-genomes-reproducibility. Cached embeddings and intermediate artefacts are available from the authors on request.

## Supplementary Material

The Supplementary Material for this article can be found online at: [Link will be provided by Frontiers].

## Notes

### Competing Interest Statement

The authors have declared no competing interest.

