## Supplementary Material for "Genomic foundation model embeddings encode higher-order viral genome architecture beyond sequence composition: a benchmark of Evo 2"

### 1 Supplementary Tables

**Supplementary Table S1.** Precision control on the NVIDIA H100. Cross-validated probe performance (repeated cluster-aware cross-validation, mean ± SD) for the Evo 2 20B embedding extracted with native FP8 input projections versus the same layers re-extracted with FP8 disabled (forced bfloat16), on identical genomes and folds. Classification reports accuracy; regression reports $R^2$. Forcing bfloat16 degrades every target, most severely the fine-grained architectural features (coding fraction, gene density, mean intergenic length), indicating that the 20B checkpoint is calibrated for FP8 inference rather than precision-agnostic. Each model is therefore reported in its native operating configuration (Section 2.5).

| Probe target | blocks.15 FP8 | blocks.15 bfloat16 | blocks.18 FP8 | blocks.18 bfloat16 |
|---|---|---|---|---|
| Baltimore class (acc) | 0.962 ± 0.007 | 0.835 ± 0.020 | 0.961 ± 0.011 | 0.830 ± 0.018 |
| Host domain (acc) | 0.992 ± 0.004 | 0.944 ± 0.012 | 0.995 ± 0.005 | 0.940 ± 0.015 |
| Family (acc) | 0.897 ± 0.023 | 0.835 ± 0.039 | 0.907 ± 0.032 | 0.854 ± 0.034 |
| coding_fraction ($R^2$) | 0.606 ± 0.069 | 0.196 ± 0.072 | 0.606 ± 0.064 | 0.240 ± 0.075 |
| gene_density ($R^2$) | 0.793 ± 0.038 | 0.417 ± 0.100 | 0.766 ± 0.033 | 0.453 ± 0.079 |
| noncoding_bp, log ($R^2$) | 0.757 ± 0.044 | 0.594 ± 0.049 | 0.752 ± 0.036 | 0.620 ± 0.042 |
| n_genes, log ($R^2$) | 0.903 ± 0.034 | 0.818 ± 0.046 | 0.908 ± 0.029 | 0.838 ± 0.041 |
| mean_intergenic_len, log ($R^2$) | 0.556 ± 0.052 | 0.273 ± 0.082 | 0.556 ± 0.050 | 0.337 ± 0.046 |

**Supplementary Table S2.** Effect of sequence-identity–aware cross-validation. Probe performance under random repeated cross-validation versus group-aware cross-validation in which MMseqs2 linclust clusters (95% identity, 85% coverage) are used as groups, so that no cluster spans training and held-out folds. Δ is the change induced by the group-aware scheme. Only 16 of the 1,912 probed genomes merged into clusters, and no metric changes by more than 0.012 in either model, confirming that the reported scores are not inflated by near-duplicate leakage.

| Probe target | 20B random CV | 20B cluster-aware CV | Δ | 7B random CV | 7B cluster-aware CV | Δ |
|---|---|---|---|---|---|---|
| Baltimore class (acc) | 0.957 ± 0.016 | 0.961 ± 0.011 | +0.004 | 0.890 ± 0.015 | 0.883 ± 0.025 | -0.007 |
| Host domain (acc) | 0.993 ± 0.006 | 0.995 ± 0.005 | +0.002 | 0.973 ± 0.007 | 0.971 ± 0.009 | -0.002 |

| Probe target | 20B random CV | 20B cluster-aware CV | Δ | 7B random CV | 7B cluster-aware CV | Δ |
|---|---|---|---|---|---|---|
| Family (acc) | 0.913 ± 0.017 | 0.907 ± 0.032 | -0.006 | 0.860 ± 0.035 | 0.857 ± 0.030 | -0.003 |
| coding_fraction ($R^2$) | 0.604 ± 0.033 | 0.606 ± 0.064 | +0.002 | 0.625 ± 0.056 | 0.618 ± 0.066 | -0.007 |
| gene_density ($R^2$) | 0.771 ± 0.036 | 0.766 ± 0.033 | -0.005 | 0.699 ± 0.047 | 0.691 ± 0.050 | -0.008 |
| noncoding_bp, log ($R^2$) | 0.745 ± 0.039 | 0.752 ± 0.036 | +0.007 | 0.692 ± 0.043 | 0.692 ± 0.029 | -0.000 |
| n_genes, log ($R^2$) | 0.898 ± 0.028 | 0.908 ± 0.029 | +0.010 | 0.823 ± 0.029 | 0.825 ± 0.039 | +0.002 |
| mean_intergenic_len, log ($R^2$) | 0.552 ± 0.034 | 0.556 ± 0.050 | +0.004 | 0.355 ± 0.079 | 0.367 ± 0.059 | +0.012 |

**Supplementary Table S3.** Dimensionality-matched comparison. Because the 20B embedding is higher-dimensional than the 7B (8,192 vs 4,096 components), which could by itself inflate linear-probe performance, every probe was re-run with both embeddings projected onto a common 150-component PCA subspace fit inside each cross-validation fold (to prevent leakage). The 20B advantage persists at matched dimensionality, indicating an effect of model scale rather than of embedding width. All values are cluster-aware cross-validation, mean ± SD.

| Probe target | 20B full (8,192-d) | 20B PCA-150 | 7B full (4,096-d) | 7B PCA-150 |
|---|---|---|---|---|
| Baltimore class (acc) | 0.961 ± 0.011 | 0.958 ± 0.009 | 0.883 ± 0.025 | 0.883 ± 0.023 |
| Host domain (acc) | 0.995 ± 0.005 | 0.991 ± 0.005 | 0.971 ± 0.009 | 0.971 ± 0.010 |
| Family (acc) | 0.907 ± 0.032 | 0.913 ± 0.031 | 0.857 ± 0.030 | 0.856 ± 0.032 |
| coding_fraction ($R^2$) | 0.606 ± 0.064 | 0.597 ± 0.063 | 0.618 ± 0.066 | 0.617 ± 0.065 |
| gene_density ($R^2$) | 0.766 ± 0.033 | 0.733 ± 0.032 | 0.691 ± 0.050 | 0.664 ± 0.045 |
| noncoding_bp, log ($R^2$) | 0.752 ± 0.036 | 0.722 ± 0.036 | 0.692 ± 0.029 | 0.665 ± 0.031 |
| n_genes, log ($R^2$) | 0.908 ± 0.029 | 0.875 ± 0.028 | 0.825 ± 0.039 | 0.803 ± 0.043 |
| mean_intergenic_len, log ($R^2$) | 0.556 ± 0.050 | 0.510 ± 0.051 | 0.367 ± 0.059 | 0.333 ± 0.048 |

**Supplementary Table S4.** Composition of the viral genome corpus (n = 19,429 RefSeq records, including individual segments of segmented viruses), cross-tabulated by Baltimore replication class and host domain as assigned through the ICTV Virus Metadata Resource (MSL41). Records whose family- or genus-level taxonomy could not be mapped to a Baltimore class or to a host domain in the VMR are reported as unassigned/unknown. Probe subsets were drawn as balanced samples from these quota groups (Section 2.4), so that the strong imbalance of the full corpus — for example, 6,311 class I records versus 94 class VI — does not propagate into the probes.

| **Baltimore class** | **Eukaryote** | **Bacteria** | **Archaea** | **Unknown** | **Total** |
|---|---|---|---|---|---|
| I (dsDNA) | 1,003 | 5,179 | 129 | 0 | 6,311 |
| II (ssDNA) | 2,283 | 132 | 17 | 0 | 2,432 |
| III (dsRNA) | 1,627 | 36 | 0 | 0 | 1,663 |
| IV (ssRNA+) | 2,715 | 834 | 0 | 0 | 3,549 |
| V (ssRNA−) | 2,005 | 0 | 0 | 0 | 2,005 |
| VI (ssRNA-RT) | 94 | 0 | 0 | 0 | 94 |
| VII (dsDNA-RT) | 137 | 0 | 0 | 0 | 137 |
| Unassigned | 712 | 0 | 0 | 2,526 | 3,238 |
| **Total** | 10,576 | 6,181 | 146 | 2,526 | 19,429 |
